# Next-generation *in situ* conservation and educational outreach in Madagascar using a mobile genetics lab

**DOI:** 10.1101/650614

**Authors:** Marina B. Blanco, Lydia K. Greene, Rachel C. Williams, Lanto Andrianandrasana, Anne D. Yoder, Peter A. Larsen

## Abstract

Madagascar is a biodiversity hotspot that is facing rapid rates of deforestation, habitat destruction and poverty. Urgent action is required to document the status of biodiversity to facilitate efficacious conservation plans. Within country, new generations of Malagasy scientists and conservationists are taking on leadership roles, although many lack access to modern genetic sequencing and are underrepresented in academic publications, when compared to international counterparts.

With the recent advent of portable and affordable genetic technologies, it is now possible to tackle logistical considerations. Mobile genetics labs, with the capacity for *in situ* DNA extraction, amplification and sequencing, can produce scientifically reproducible data under field conditions, minimizing the time between sample collection and data analysis. Additionally, mobile labs offer powerful training opportunities for in-country scientists that enable local students and researchers to actively participate and contribute fully to the research enterprise, and that further empower these communities to contribute to the conservation dialog.

Here, we show “proof of concept” by deploying a miniaturized thermal cycler alongside the Oxford Nanopore MinION DNA sequencer in Madagascar, including in the newly established Anjajavy Protected Area in northwestern Madagascar. We successfully extracted DNA from tissue samples collected using minimally-invasive techniques, amplified and sequenced a phylogenetically informative mitochondrial gene (cytochrome-b; *cytb*), and thereby confirmed the presence of Danfoss’ mouse lemur (*M. danfossi*) within the Anjajavy Reserve.

To demonstrate the reproducibility of our methods, we successfully performed our established molecular and analytical pipeline at two additional locations in Madagascar, where we also conducted two-day workshops at local higher-education Institutions to demonstrate the process from tissue samples to DNA sequencing. Ultimately, we show that a mobile genetics lab can provide reliable and expeditious results, become a powerful educational tool, and allow scientists to conduct genetic analyses, potentially allowing for rapid interventions under emergency conditions *in situ*.

## Introduction

Madagascar, one of the world’s most threatened biodiversity hotspots, is fighting severe challenges to the long-term survival of its endemic wildlife due to habitat loss and degradation, while experiencing one of the fastest population growth rates worldwide (Gardner et al. 2018; UNFPA 2018). Thus, there is a sense of urgency to accurately report biodiversity to assess conservation risks and translate these data into policy action. Somewhat paradoxically, thanks to advances in genetic technologies, our quantification of biodiversity levels continues to rise despite high deforestation rates. This is largely the result of new biological surveys targeting poorly known areas, the integration of genetic analyses with more traditional morphological assessments, and a barcoding approach to species identification. Oftentimes, as new species are formally described in the literature, they are immediately tagged as “endangered” or “critically endangered” under International Union for Conservation of Nature (IUCN) regulations, because their distributions are restricted, and their habitats are highly fragmented.

This is particularly true of the lemurs of Madagascar, which are currently considered the most threatened group of primates on earth (Estrada et al. 2017). Among them, the “cryptic” nocturnal mouse lemurs (*Microcebus*) have undergone one of the most dramatic taxonomic expansions, with species numbers increasing from only a few to 24 in the last decades (Hotaling et al. 2016). Some mouse lemur species are known to live in sympatry and, in certain cases, there is evidence of hybridization between them, which make species assignation by phenotypic cues challenging at best (Hapke et al. 2011). Further increasing their appeal as research models, genomic data are rapidly accruing, including a genome assembly to chromosome-level for the gray mouse lemur (*Microcebus murinus*) (Larsen et al. 2017).

On the flip side of its biodiversity wealth, Madagascar’s academic opportunities are rare and limiting. Species descriptions and updates on the conservation status of lemurs have been traditionally led by foreign researchers. Although there is an ongoing trend for Malagasy scientists to take a more active role in project design, data collection and analysis, international collaborations are still vital and encouraged in Madagascar, both to contribute financially and technologically, and to facilitate knowledge production and dissemination. Efforts by Malagasy researchers to encourage these collaborations are especially laudable in the context of national policy and education. Unfortunately, reliance on international support has led foreign researchers to take leadership in publishing and securing funding. This is evident by the underrepresentation of lead-authors affiliated with Malagasy institutions, a trend expected to contribute to weakening high education development and quality for years to come (Waeber et al. 2016).

Yet, we are now at scientific, academic, and technological crossroads: A new generation of Malagasy researchers are establishing labs and/or developing research programs in country. At the same time, new DNA sequencing technologies are revolutionizing the fields of genetics and genomics, expanding applications worldwide through the creation of miniaturized devices. These new technological products are relatively affordable and have the potential to sequence even whole genomes in real time (Tyler et al. 2018). At the forefront of these developments are devices released from Oxford Nanopore Technologies (ONT) and miniPCR. These companies have created the MinION (ONT; a portable nanopore-based DNA sequencer platform) and the miniPCR (a miniaturized thermal cycler). These technologies have already been tested under field conditions in a variety of projects around the world, including rapid species assessments in a South American biodiversity hotspot (Pomerantz et al. 2018), disease surveillance to cope with epidemiological crises in West Africa (Quick et al. 2016) characterization of microbiome communities under extreme climatic conditions such as Antarctica (Johnson et al. 2017), and even in outer space (Castro-Wallace et al. 2017). As these technologies are becoming widely available, there is increasing need for operational workflows to simplify analyses and reduce laboratory costs, while being able to survey remote field sites and produce expedited and reliable results (Maestri et al. 2019). And far from least, these technologies are proving to be excellent platforms for engaging in-country students and scientists in the fast-moving area of field genomics (Watsa et al. 2019).

Our interdisciplinary research team has been conducting research in Madagascar for decades. Recently, we have furnished a mobile genetics lab for the field, both for research and capacity building. Our objectives were twofold: 1-to test the efficacy of a mobile genetics lab in Madagascar by sequencing a marker gene from mouse lemurs (i.e., genetic “barcoding”) and providing the first lemur species assessment *in situ*; 2- to build in-country capacity by conducting workshops at local academic Institutions in Madagascar to gauge the interest and the potential use of the lab as an educational tool for high education students.

## Methods

### Lab implementation in Madagascar

Our mobile genetics lab included equipment, reagents and supplies needed to process samples from DNA extraction to sequencing (Table 1). We tested all lab components at Duke University, North Carolina, USA. The lab was fitted in two Pelican cases and shipped from the USA to Madagascar in May 2018 (Sup. Inf.). In Sambava, NE Madagascar, we conducted our first lab test in country, checking reagents and flow cells, testing equipment and troubleshooting lab protocols to local conditions. This comprehensive testing included DNA extractions, PCRs, library preparations and sequencing, using tissue samples stored at the office from previous research missions. Moreover, Moreover, MBB and LKG trained LA in the basics of lab operation. Then, the three of us organized a two-day workshop conducted at the Centre Universitaire Régional de la SAVA (CURSA) a local branch of the University of Antsiranana in northeastern Madagascar.

**Table 1.**
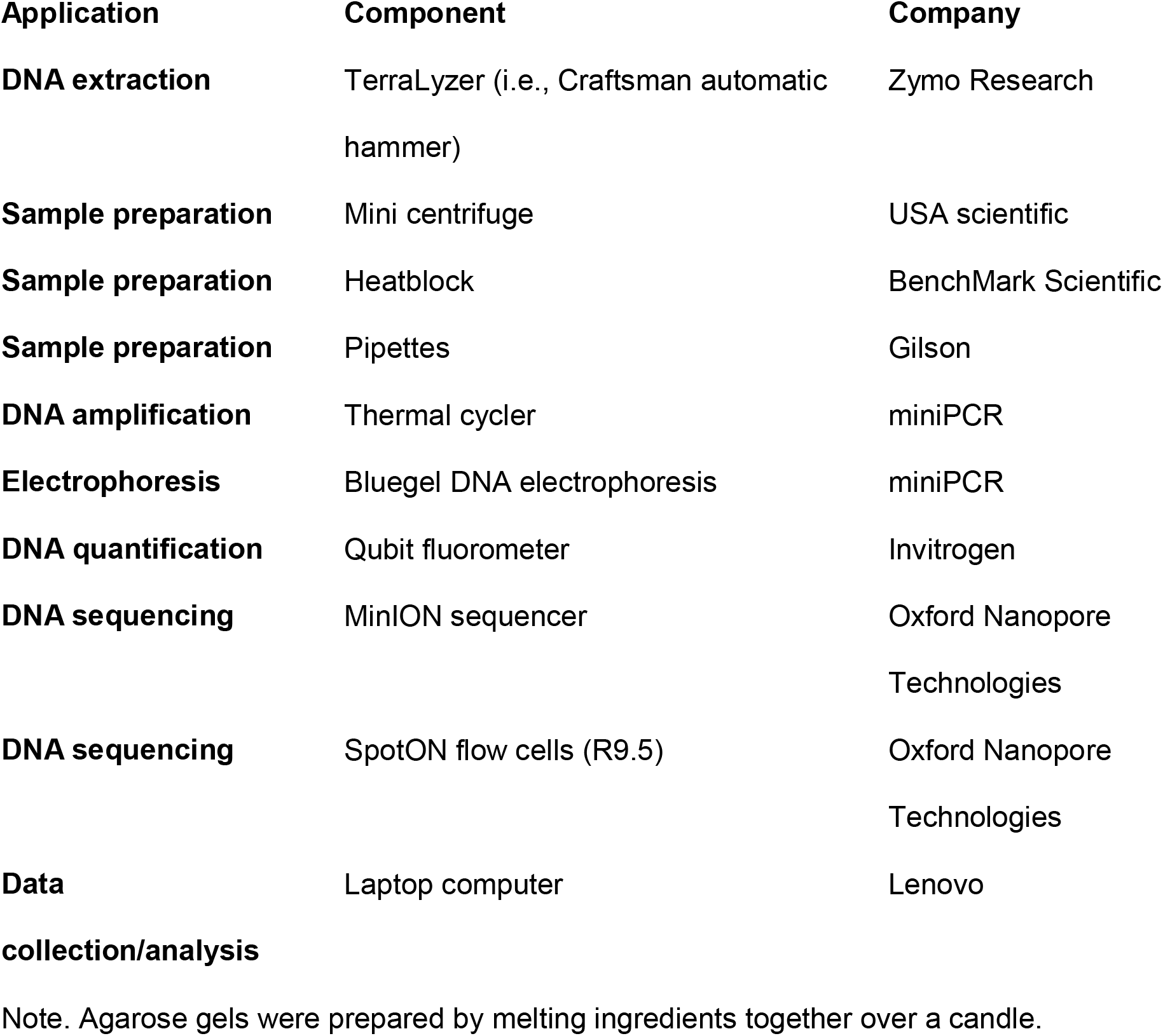
Mobile genetics lab components

| **Application** | **Component** | **Company** |
| --- | --- | --- |
| **DNA extraction** | TerraLyzer (i.e., Craftsman automatic hammer) | Zymo Research |
| **Sample preparation** | Mini centrifuge | USA scientific |
| **Sample preparation** | Heatblock | BenchMark Scientific |
| **Sample preparation** | Pipettes | Gilson |
| **DNA amplification** | Thermal cycler | miniPCR |
| **Electrophoresis** | Bluegel DNA electrophoresis | miniPCR |
| **DNA quantification** | Qubit fluorometer | Invitrogen |
| **DNA sequencing** | MinION sequencer | Oxford Nanopore Technologies |
| **DNA sequencing** | SpotON flow cells (R9.5) | Oxford Nanopore Technologies |
| **Data collection/analysis** | Laptop computer | Lenovo |

Note. Agarose gels were prepared by melting ingredients together over a candle.

In early July, MBB and LKG moved the lab from Sambava to a field site, the Anjajavy Lodge and Reserve. This site was ideal because it has electricity, an unusually high density of mouse lemurs and, importantly, because the species of mouse lemur at this site has/have yet to be genetically confirmed. Anjajavy (S14.99025 E47.22958) encompasses ~1,200 ha of private Reserve, and ~8,000 ha of recently protected area, predominantly dry deciduous forest. We conducted our field mission between July 9 and 29, 2018. During the first 10 days of our mission, we captured and obtained tissue samples from 12 mouse lemurs. During the last 10 days in the field, we extracted all tissue samples and chose a high-quality extraction to sequence. The process from DNA extraction until sequencing took ~ 8 hours and was partitioned over two days (Sup. Inf.).

After Anjajavy, we traveled to the capital Antananarivo to set the lab at the Vahatra Association office, the leading Malagasy NGO promoting capacity building and scientific research in Madagascar. With logistical assistance from Vahatra staff, we conducted our second workshop at the Vahatra Association’s office, for university students and researchers interested in genetics.

### Sample processing and analysis

Genomic DNA was extracted from tissue samples using DNeasy Blood & Tissue Kit (Qiagen, Hilden, Germany) according to the manufacturer’s protocol. We incorporated mechanical lysis (i.e., bead beating) as an additional step after chemical lysis. We selected to sequence the entirety of the mitochondrial gene, *cytb*, (~1100bp) which has proven a reliable phylogenetic marker for mouse lemurs (Hotaling et al. 2016). We amplified c*ytb* using the miniPCR thermal cycler. We used a PCR cleanup protocol with AMPure beads before proceeding to sequencing. DNA library preparation and flow cell loading were carried out according to the protocols by ONT. Flow cell pore availability ranged between 85 and >1000 active pores (Sup. Inf.).

We allowed each library to run for 1 hour, and we used MinKNOW software to base call for 3-8% of bases. This low percentage was enough to accomplish great depth in sequencing coverage with more than 10,000X of the *cytb* amplicon per sample. Sequenced data were stored as FASTq files and retrieved using MinKNOW software. Raw reads were first filtered by read length between 1000 and 1400 bases (approximating that of the *cytb* gene) and then mapped to a *M. murinus cytb* reference sequence (GenBank accession number: U53572) before generating a consensus sequence. Consensus sequences were aligned to the reference sequence using Geneious software, and then blasted to our database comprising ~ 270 published (NCBI GenBank) and unpublished mouse lemur sequences from the Yoder lab. Finally, we created a phylogenetic tree using Neighbor-Joining (NJ) analysis with uncorrected p-distances as implemented in PAUP version 4a165. One thousand bootstrap trees were also estimated by NJ with PAUP and we used RAxML version 8.2.12 to draw bootstrap support onto the nodes from the original data set. The phylogenetic tree was edited in FigTree version 1.4.4.

All research protocols complied with Institutional Animal Care and Use of Animals at Duke University (IACUC# A263-17-12), and field research was approved by the Ministry of Environment, Ecology and Forests of Madagascar (Permit# 035/18/MEEF/SG/DGF/DSAP/SCB.Re).

## Results

In total, we sequenced DNA from four mouse lemurs using the mobile genetics lab in Madagascar. Two mouse lemur samples were sequenced in Sambava, one sample was sequenced *in situ* at the Anjajavy field site, and one sample was sequenced in Antananarivo (Fig. 1). All mouse lemur samples grouped with those of *Microcebus danfossi*, thus we genetically confirmed the presence of this species at Anjajavy (Fig. 2). Consensus sequences generated in this study were stored in GenBank under accession numbers XXX-XXX.

**Fig. 1.**
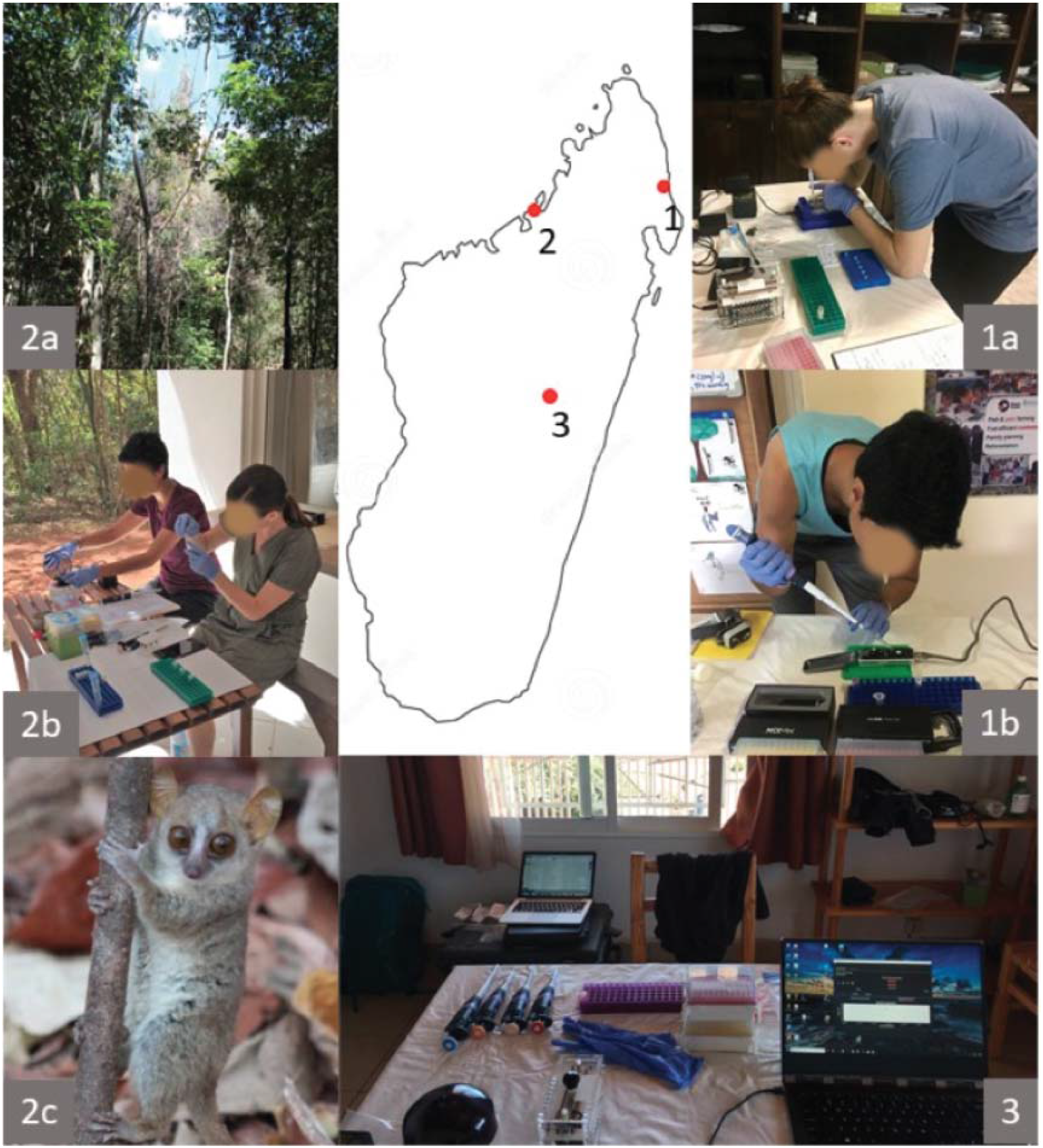
Locations for the use of the genetics lab. 1.Sambava: (a) Loading gel in the DLC/SAVA office, (b) Loading MinION prior to sequencing; 2. Anjajavy: (a) deciduous forest, (b) lab setting near forest, (c) Danfoss’ mouse lemur; 3. Antananarivo: lab setting at Vahatra office.

**Fig. 2.**
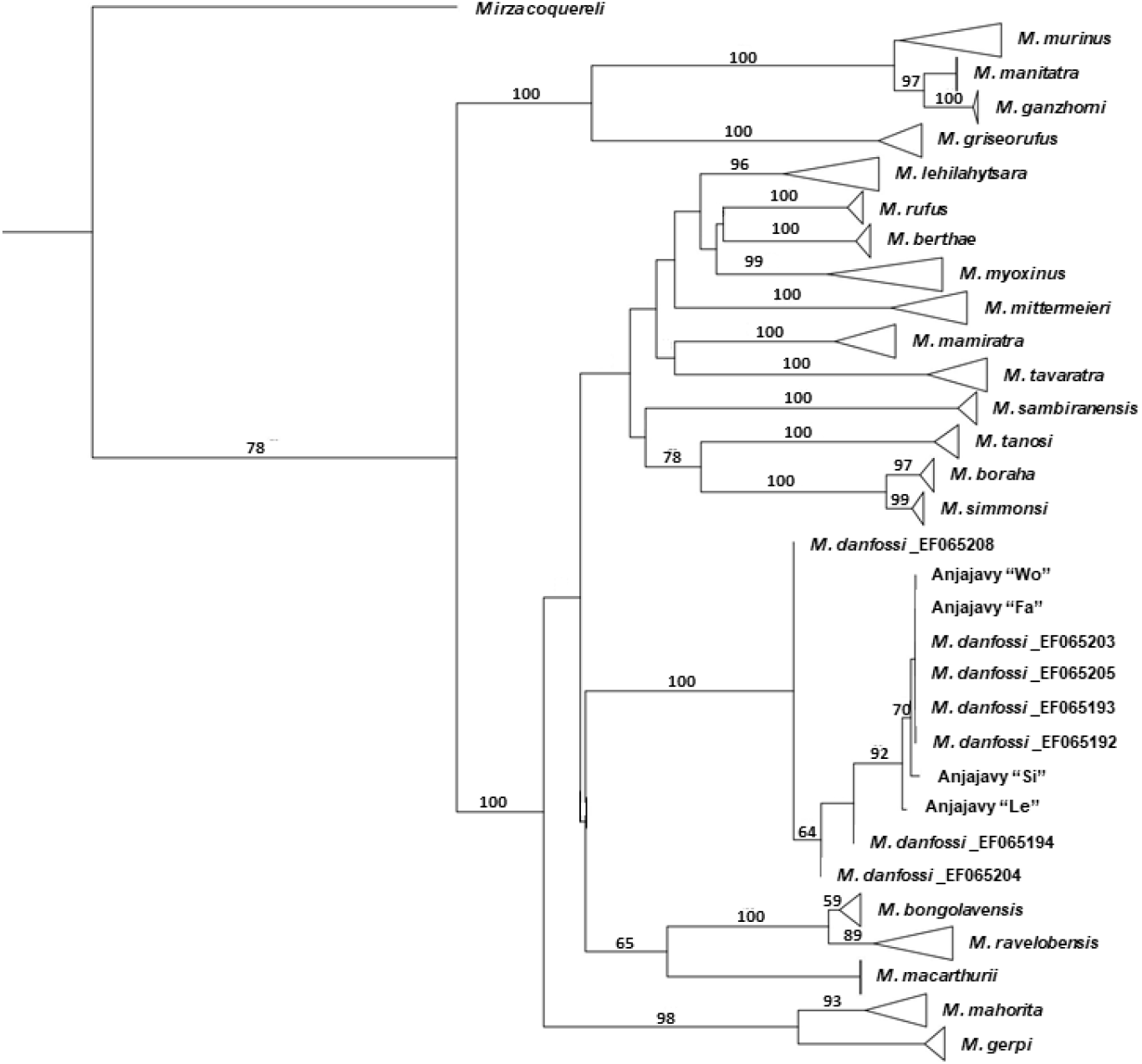
*Microcebus* phylogeny based on *cytb* sequences. Branches were collapsed for practical purposes, except for the *M. danfossi* lineage, to highlight consensus sequences from Anjajavy; node support above 50 % is shown. GenBank accession numbers for Anjajavy samples are pending, those for published sequences of Danfoss’ mouse lemurs are listed next to the species name.

We conducted two workshops to target college students, at the Centre Universitaire Régional de la SAVA (CURSA) in the town of Antalaha, and at the Vahatra Association office, in the capital Antananarivo (Fig. 3). During both workshops, we described and demonstrated all procedures from tissue extraction to sequencing to species assignation. A total of 66 students attended the workshop at CURSA conducted in Malagasy, where some students actively participated in hands-on molecular techniques such as pipetting, loading samples in the miniPCR thermal cycler, setting the agarose gel and loading samples using the Bluegel DNA electrophoresis kit. At Vahatra Association, a total of 25 students (the maximum room capacity) attended the workshop, which was conducted in English. These participants had more comprehensive backgrounds in genetics, so workshop participants engaged in all lab activities as well as discussions about potential implementation of this technology in a variety of research topics. During the workshop at Vahatra Association, we sequenced a mouse lemur sample from our recent field mission at Anjajavy.

**Fig. 3.**
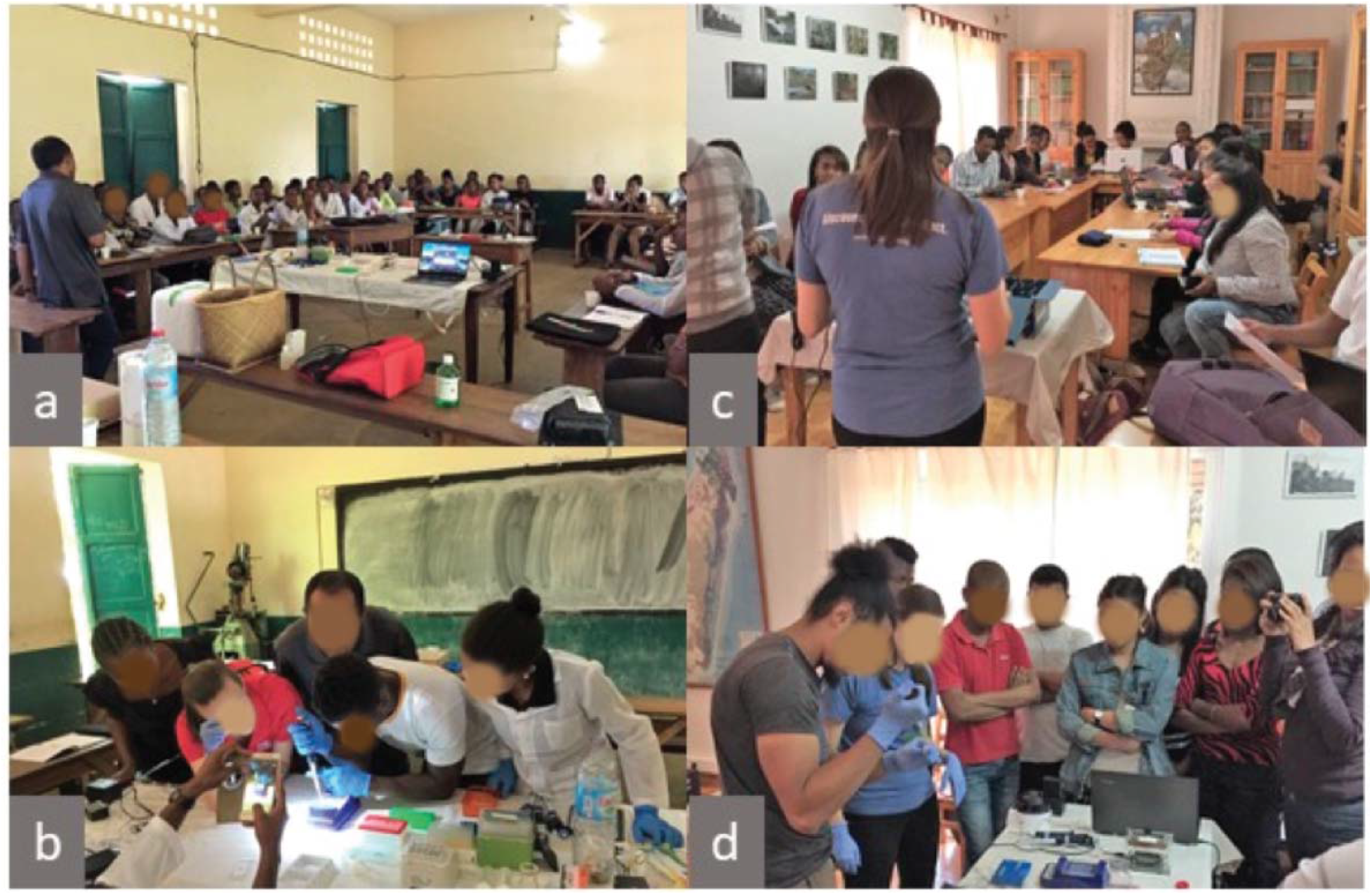
Workshops in Madagascar: a,b at CURSA in Antalaha; c,d at Vahatra office in Antananarivo

## Discussion

Portable and affordable technologies in the field of genetics have made it feasible to produce scientifically robust data under field conditions, minimizing the time between sample collection and data analysis. The capacity for *in situ* genetics also provides training opportunities to enable active participation of local students in the whole process of scientific research.

To our knowledge, we were the first researchers to perform *in situ* sequencing of wild lemurs in Madagascar. We were able to confirm species assignment for one of the lemurs at Anjajavy within a week, from lemur capture to tissue collection, to the generation of a phylogenetic tree. Our results are consistent with the known distribution of *M. danfossi*, between the Sofia and Maevarano rivers, NW Madagascar (Olivieri et al. 2007). Anjajavy’s mouse lemurs had been tentatively assigned as *M. danfossi* on the basis of morphological assessments (Randrianambinina et al. 2010), but we provide the first genetic confirmation of Danfoss’ mouse lemurs at this site.

We would like to emphasize that mobile genetics technology not only minimizes time between sample collection and analysis, but has the potential to address urgent conservation concerns in *real time*. Conservation crises can span from needing rapid biodiversity assessments in threatened habitats, to dealing with wildlife die-off situations or disease-related cases that require immediate intervention (e.g., Carver 2018).

We showed “proof of concept” that this technology can be deployed in remote sites, that results can be obtained in a speedy manner and that training sessions can prepare advanced students with the skills and means to conduct genetic analysis *in situ*. As a testament to the latter point, both MBB and LKG who deployed the lab in Madagascar, had some experience with molecular genetics and analysis, but are not geneticists themselves. Week-long training in the USA provided the necessary skills, tools and information to run the lab on the ground in Madagascar and train LA and others in its use.

One unexpected corollary of the workshops was the general interest from participants to apply these technologies to a large range of research topics. We also learned that there are facilities and Institutions in country already using MinION technology such the Mahaliana Lab (https://www.mahaliana.org/) and the Pasteur Institute (http://www.pasteur.mg/). Connectivity among the scientific community, both national and international, will be beneficial for researchers interested in funding, contracting services or collaborating with them. Finally, we showed that the mobile genetics lab can be a powerful educational tool, for teaching basic concepts to introductory students, or exemplifying complex procedures for advanced college students with background in biological sciences. Thus, the potential for this technology to immediately produce data, coupled with the power of capacity building to engage local researchers significantly outweighs the logistical challenges to obtain, transport and maintain lab supplies and reagents in remote settings.

In sum, miniaturized and more affordable technologies have the potential not only to speed up production of knowledge and solve biological and environmental crises in efficacious manners at remote settings, but also to shape the professional careers of passionate scientists in less advantageous academic settings and to level the scientific playing field.

## Acknowledgements

We thank the Malagasy Government for authorizing us to conduct research in Madagascar. We thank A. Raselimanana for his help advertising the workshop at CURSA, Antalaha, and A. Raselimanana for organizing the workshop at Vahatra office, Antananarivo. We also thank S. Goodman for allowing us to use the Vahatra office for the workshop. Equipment and supplies for the mobile lab were made possible by a private donation to PAL and the Duke Lemur Center from W. Korman and Google Inc. Funding for this project was provided by Global Wildlife Conservation’s Lemur Conservation Action Fund and IUCN SOS. Funding for LKG was provided by an NSF DDRIG BCS 1749898. Additional travel and supplies were supported by funds from the John Simon Guggenheim Foundation to ADY. We thank S. Bornbusch, R. Schopler and K. Thompson for acquiring and transporting supplies to Madagascar. We are also grateful to C. de Foucault, E. Rambeloson, H. Rasoanaivo and Anjajavy staff members for their assistance in the field. We are grateful to G. Tiley for his assistance with the phylogenetic analysis. This is a DLC publication # XXX.

## Supporting Information

Travel itinerary, Laboratory protocols, and Sampling methods. The authors are solely responsible for the content and functionality of these materials. Queries (other than absence of the material) should be directed to the corresponding author.

## Supporting Information

### Travel itinerary

Our genetics lab was shipped in two Pelican cases from Washington Dulles Airport, USA to Ivato Airport Antananarivo, Madagascar in early May. The trip included two flights and was ~ 27 hours total. The following morning, the lab was shipped to the town of Sambava (NE Madagascar) via air freight, and less than 24 hours later, it was brought by car to the Duke Lemur Center/SAVA Conservation office for storage with an available fridge/freezer unit. In between international and domestic flights, sensitive reagents and flow cells were kept in coolers and refrigerated with ice packs. In early July, the lab was shipped back from Sambava to Antananarivo via air freight and, less than 48 hours later, we rented a car and drove to the town of Mahajanga, (NW Madagascar) for 13 hours. Less than 24 hours later, we rode a boat from Mahajanga to the field site, the Anjajavy Lodge and Reserve, a trip that lasted 4.5 hours. In between transportation routes, we placed reagents and flow cells under refrigeration and replaced ice packs. In late July, we reverted the original route, taking the genetics lab from Anjajavy back to Antananarivo.

### Lab protocols

#### Primers and PCR

Primers to amplify *cytb* gene were L14724: 5′–CGA AGC TTG ATA TGA AAA ACC ATC GTT G–3′, and H15915: 5′–AAC TGC AGT CAT CTC CGG TTT ACA AGA C–3 as previously described in Irwin et al. (1991). Each Polymerase chain reaction (PCR) contained approximately 1.5 *μ*L of PCR product, 12.5 *μ*L LongAmp Taq DNA Polymerase (New England Bio Labs), 1.25 *μ*L of each primer, and 8.5uL water for a 25 *μ*L total volume. Samples for the PCR run followed the following settings: initial denaturation 95°C for 2 minutes, 32 cycles of denaturation at 95°C for 30 seconds, annealing at 57°C for 30 seconds, extension at 72°C for 45 seconds, and a final extension at 72°C for 480 seconds. PCR products were then cleanup using AMPure beads and washed in 70% EtOH before resuspending in water.

#### Library preparation

DNA products (45 μL amplicon) were prepared with 7 μL Ultra II End-prep reaction buffer and 3 μL Ultra II End-prep enzyme and 5 μL water. Samples were washed using AMPure beads and 70% EtOH and resuspended in 30 μL water. Adapter ligation and tethering was then carried out with 20 *μ*L of Adapter Mix (ONT) and 50 *μ*L of NEB Blunt/TA ligation Master Mix (New England Biolabs). The adapter ligated DNA library was then purified with AMPure beads and ABB buffer. Samples were resuspended in 13 μL water and placed in a LoBind tube ready for sequencing.

#### Sequencing

We prepared pre sequence-library following ONT protocols, by mixing 12 μL amplicon, 2.5 μL water, 25.5 μL LBB and 35 μL RBF. We inserted flow cell in the MinION frame, and loaded the flow cell’s priming port with 800 μL of priming mix (576 μL RBF and 624 μL water) with SpotON cover closed. We additionally loaded 200 μL priming mix in priming port with SpotON cover open. The sample was then added to SpotOn por via dropwise fashion. Finally, we covered SpotOn and priming ports, close the MinION lid and open the MinKNOW GUI software to proceed to sequence. Note: We were authorized by ONT to use MinKNOW 18.5.1.0 version to base call offline, because we were off grid at Anjajavy.

#### Flow cells

We used a total of three flow cells (R 9.4), one at each location: Sambava, Anjajavy and Antananarivo respectively. Flow cell number one, used in Sambava, was transferred with the mobile genetics lab in May, and contained more than 1000 active pores at the time of use, well over the 800 threshold recommended by company. Flow cell number two was brought by a researcher to Anjajavy in July, and contained 85 active pores. Despite the low count, likely the result of disrupted transportation conditions, results obtained from this unit were reliable. In fact, our ability to reliably sequence *cytb* gene speaks to the great redundancy buffer provided by flow cells, when relatively short and single genes are the source of sequencing. Flow cell number three was brought by another researcher in September to Antananarivo, and used for our last sequencing event. This flow cell contained over 1000 active pores at the time of sequencing.

#### Note

When we tested the lab at Duke University, we sequenced a sample from a grey mouse lemur (*M. murinus*) from the Duke Lemur Center using the MinION sequencer, and generated a consensus sequence that was 100% identical to the reference sequence that had been generated by Sanger sequencing.

### Sampling methods

Mouse lemurs were captured with Sherman traps (3 x 3.5 x 9”) baited with small pieces of banana and set along trails at 1.5m height. Traps were set in the afternoon and checked in the mornings for 6 days. All captured lemurs were brought back to the campsite for processing. At the campsite, individuals were weighed, measured, and microchipped for identification (Trovan®). Tissue samples (ear biopsies, 2mm) were taking from anesthetized individuals (Ketamine, 10mg/kg) and stored in 90% alcohol for further analysis. Individuals were released at trapping sites later the same day.

